# Predictive design of tissue-specific mammalian enhancers that function *in vivo* in the mouse embryo

**DOI:** 10.64898/2025.12.22.695948

**Authors:** Shenzhi Chen, Vincent Loubiere, Ethan W. Hollingsworth, Sandra H. Jacinto, Atrin Dizehchi, Jacob Schreiber, Evgeny Z. Kvon, Alexander Stark

## Abstract

Enhancers control tissue-specific gene expression across metazoans. Although deep learning has enabled enhancer prediction and design in mammalian cell lines and invertebrate systems, it remains unclear whether such approaches can operate within the regulatory complexity of mammalian tissues in vivo. Here, we present a general strategy for designing tissue-specific enhancers that function reliably in mice. We use deep learning to train compact convolutional neural networks (CNNs) on genome-wide chromatin accessibility and fine-tune them via transfer learning on validated human and mouse enhancers. Guided by these models, we design fifteen synthetic enhancers for the heart, limb, and central nervous system (CNS) in mouse embryos, all of which are active in their intended target tissue. Our work establishes a generalizable framework for programmable control of mammalian gene expression *in vivo*, opening new avenues in functional genomics, synthetic biology, and gene therapy.

## Introduction

Enhancers are key regulatory elements that direct tissue-specific gene transcription and are widely used to drive transgene expression in defined cellular contexts^1^. Enhancer activity is known to rely on transcription factor (TF) motifs, but their precise syntax, such as motif type, number, spacing, and orientation, remains poorly understood, challenging the computational prediction and *de novo* design of synthetic enhancers. Deep learning^2,3^ has led to breakthroughs in predicting and designing synthetic enhancers^4–8^ (reviewed in ref. ^9^): Models trained on Massively Parallel Reporter Assays (MPRAs) that measure enhancer activity for thousands of enhancer candidates in cultured cells^10,11^ (reviewed in ref. ^12^) can accurately predict quantitative enhancer activity and cell-type specificity, and have been used to design synthetic enhancers that function in *Drosophila* and mammalian cell lines^4,6–8^. Although some MPRA-derived synthetic enhancers are active in vivo^7^, MPRA-based models remain limited by their reliance on cultured cells, which do not capture the spectrum of mammalian tissues. As a consequence, it is unknown whether sequence-based models can generalize to the substantially greater regulatory complexity of mammalian tissues *in vivo*.

We hypothesized that integrating genome-wide chromatin accessibility with functionally validated enhancers through transfer learning might provide the information necessary for predictive enhancer design in mammals. Indeed, our recent work in *Drosophila* embryos demonstrated that pre-training on chromatin accessibility and fine-tuning on enhancer activity can support *de novo* enhancer design in an invertebrate system *in vivo*^5^, but whether such an approach can succeed in mammals, where enhancer architectures and gene regulation are more complex and context dependent, has remained an open question.

Here we demonstrate that compact convolutional neural networks (CNNs) pre-trained on ATAC-seq data and fine-tuned on a limited set of enhancers validated by *in vivo* reporter assays in mice (VISTA Enhancer Browser^13^, Figure 1A), are sufficient to enable robust *de novo* design of mammalian enhancers that function *in vivo*. We developed sequence-to-accessibility and sequence-to-activity models for the mouse heart, limb, and central nervous system (CNS) at embryonic day 11.5 (E11.5). Guided by these models, we designed five enhancers for each tissue and tested their activities using a site-specific reporter assay in mouse embryos. All fifteen (100%) were active in their intended target tissues. These results demonstrate that mammalian enhancer function can be reliably inferred from DNA sequence alone, enabling the predictive *de novo* design of tissue-specific synthetic enhancers from modest training sets and establishing a general framework for enhancer design in mammals.

**Figure 1.**
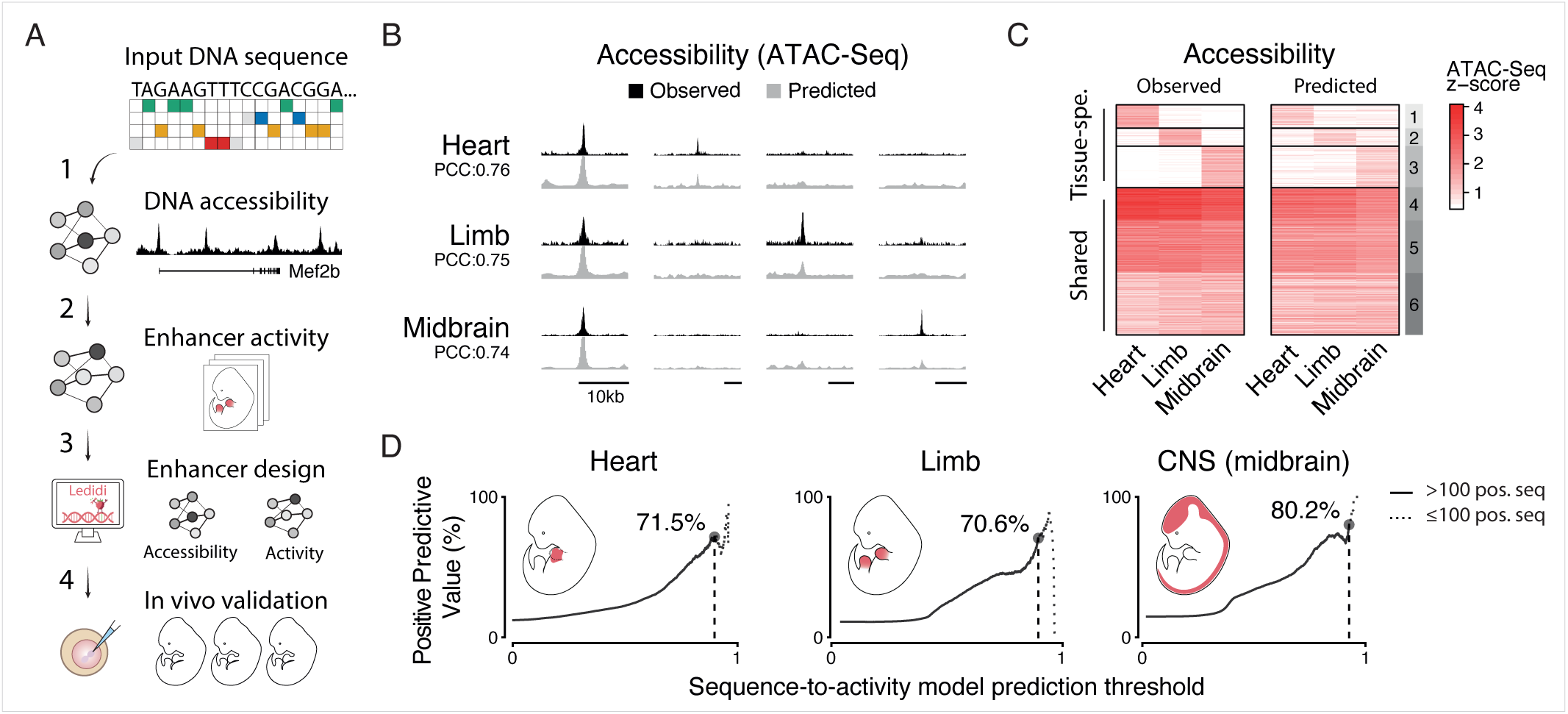
Deep learning-based prediction and design of tissue-specific mammalian enhancers (A) Schematic overview of the deep- and transfer-learning approach to enhancer design in mammals. 1) Deep-learning models are pre-trained on genome-wide DNA accessibility data before being 2) fine-tuned using transfer learning on a subset of tissue-specific enhancers from the VISTA Enhancer Browser^13^. 3) Both models are used to guide the design of synthetic enhancers using Ledidi^22^ (3), before assessing their activity and specificity in E11.5 mouse embryos using reporter assays^23^ (4). (B) Screenshot of experimentally observed (black) and predicted (grey) ATAC-seq signals at one shared and 3 tissue-specific ATAC-seq peaks on mouse chromosome 18, which was not used for training. (C) Scaled experimentally observed (left) and predicted (right) ATAC-seq signals for three tissues (shades of red, see colorbar) on held-out genomic regions that were not used for training. K-means clusters are shown on the right (k= 6). Clusters 4-6 are accessible in all three tissues. (D) PPV of enhancer activity predictions across prediction score thresholds. For each threshold (x-axis, 0–1), the y-axis shows the percentage of active enhancers (positives) among all predicted enhancers (predicted score ≥ threshold). The maximum value is reported when at least 100 positive sequences remain (dashed line).

## Results

### Deep learning-based prediction and design of tissue-specific mammalian enhancers

We first trained sequence-to-accessibility models on published ATAC-seq datasets^14^ for E11.5 mouse heart, limb and midbrain tissues. These models accurately predicted DNA accessibility on a chromosome excluded from training (mouse chr18, PCC ≥ 0.76 for each tissue; Figure S1B) and captured both shared and tissue-specific ATAC-seq peaks (Figures 1C and S1A). Similar performance was observed in cross-validation using a held-out test set of regions distributed across all chromosomes and balanced like the training set (PCC ≥ 0.88 for each tissue, Figure S1B). Notably, pairwise comparisons of experimental and predicted data between different tissues demonstrated robust prediction of differential chromatin accessibility, i.e. tissue-specific peaks (PCC ≥ 0.66; Figure S1C), making these models strong starting points to derive enhancer-activity rules.

We next fine-tuned these sequence-to-accessibility models using experimentally validated embryonic mouse and human enhancers from the VISTA Enhancer Browser through the process of transfer-learning^5,15,16^. As a design-oriented study, we prioritized precision to minimize false positives among the small number of synthetic sequences that can be tested using low-throughput, resource-intensive transgenic reporter assays *in vivo*. We therefore evaluated model performance using the Positive Predictive Value (PPV), which should reflect the expected success rate among the designed enhancers. Despite the limited number of validated tissue-specific enhancers (between 311 and 432 per tissue, Figure S1D), the resulting sequence-to-activity models achieved high PPVs, reaching 70.6% and above for enhancer sequences not used during training (neither pre-training nor transfer learning).

Notably, 74.5% of all VISTA enhancers active in the midbrain are also active in at least one other CNS subregion such as forebrain, hindbrain or neural tube, and 22.7% are active in the entire CNS (*pan-CNS*; Figure S1E). Consequently, enhancers of all other CNS sub-regions scored highly by the midbrain model, with pan-CNS enhancers reaching maximum scores (Figures S1F-G). These results suggest that the midbrain model can be used to design pan-CNS enhancers.

In addition to reaching high PPVs on the VISTA-derived test set, the fine-tuned heart, limb and CNS models successfully reject two additional sets of control sequences, 300,000 randomly generated sequences as well as 300,000 randomly sampled inaccessible genomic sequences (Figure S1H). These results suggest that the models might be able to guide *in silico* enhancer design with a high likelihood of success. Notably, models trained only on ATAC-seq data or only on validated enhancers (direct training without transfer learning) performed worse, with PPVs dropping between 20.9% and 52.1% depending on the tissue (Figures S1I-J), underscoring the value of a two-step transfer-learning approach.

Furthermore, transfer learning changed the contribution scores of various motifs, reflecting shifts in importance between sequence-to-accessibility versus sequence-to-activity models. A subset of motifs shows similarly high – or even increased – contribution scores after transfer learning, while others show decreased scores, and these changes group motifs into four clusters (Figure 2A). Specifically, motifs with maintained or increased importance (clusters 1-3) correspond to TFs enriched for developmental GO terms of the respective tissue and are preferentially expressed in that tissue^17^ (Figures S2A-B); including the master regulators MEF2 for heart^18^, TWIST1 for limb^19^, and SOX3 for CNS^20^. By contrast, motifs in a fourth cluster decrease in importance across all tissues after transfer learning and their cognate TFs are significantly enriched for broadly expressed TFs without preference for any of the three tissues (Figures S2A-B); such as CTCF that primarily functions at insulators^21^. These trends further support the role of transfer-learning in fine-tuning sequence-to-accessibility models towards predicting enhancer activity.

**Figure 2.**
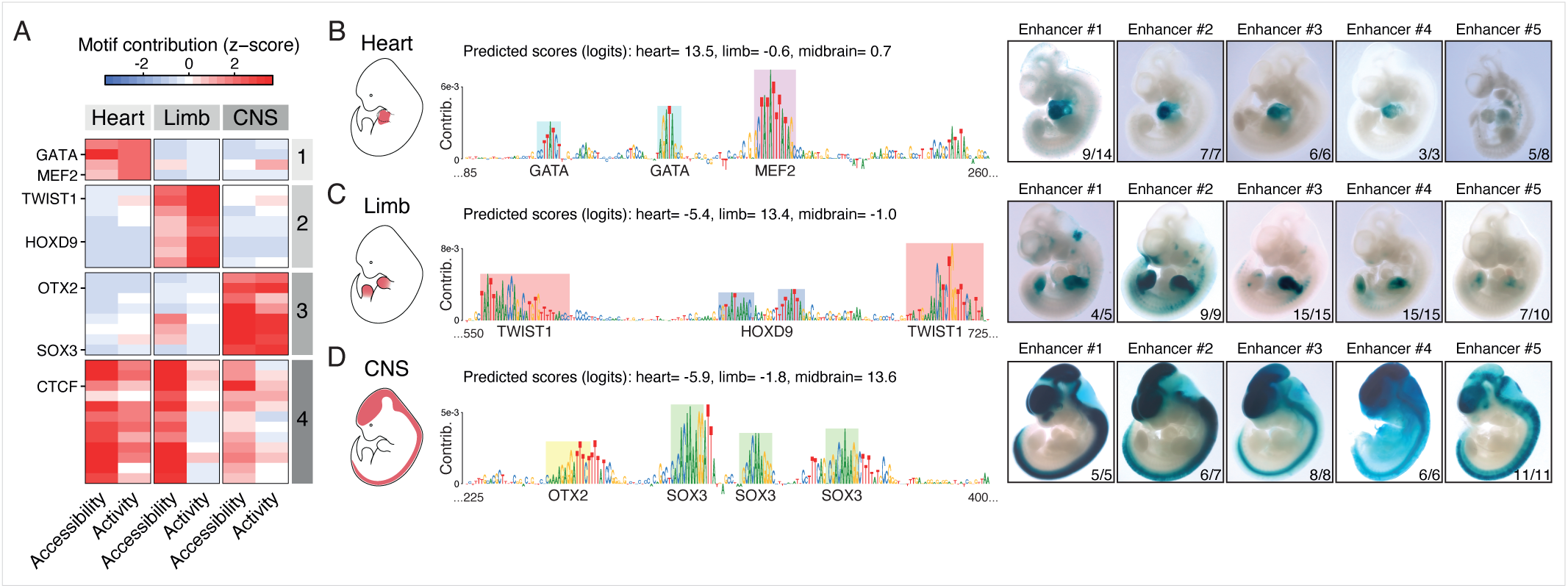
In vivo validation of tissue-specific synthetic enhancers in mouse embryos (A) Scaled motif contribution scores before (sequence-to-accessibility) and after (sequence-to-activity) transfer-learning (see x-axis; shades of red and blue, see colorbar). (B-D) Left panel: schematic view of E11.5 mouse embryos with heart (B), limb (C), and central nervous system (CNS) (D) highlighted in red. Center panel: nucleotide contribution scores at representative regions (x-axis) of designed synthetic enhancers (total length = 1,001 bp), with key TF motifs highlighted as colored boxes. Right panel: representative LacZ-stained transgenic E11.5 mouse embryos carrying five distinct synthetic enhancer sequences. The leftmost image (Enhancer #1) corresponds to the sequences illustrated in the center panel. For each enhancer, the number of embryos showing detectable activity in the target tissue is shown. Following VISTA convention^13^, synthetic enhancers that show consistent reporter gene expression among at least three embryos are positive; all embryos are shown in Figure S3.

Altogether, these results prompted us to design synthetic enhancers *de novo* for all three tissues using Ledidi, a gradient-based, model-guided sequence design framework^22^. We used both sequence-to-accessibility and sequence-to-activity models to jointly guide the design of sequences with high predicted accessibility and activity for each tissue. For each of the three tissues, the majority of the designed enhancers with high scores in their target tissue had low scores in the two other tissues (≥69.7%) and were deemed likely tissue specific (Figures S2C-D).

### In vivo validation of tissue-specific synthetic enhancers in mouse embryos

We selected five diverse candidate sequences designed to be active in the limb, heart, and CNS (15 total, Table S1). Importantly, these synthetic enhancers did not share any significant sequence similarity to the mouse or human genomes (BLAST E-value > 0.05). We then assessed their enhancer activities in E11.5 mouse embryos using our previously published site-specific transgenic reporter method, which has no background activity^23^. All fifteen synthetic enhancers are reproducibly active in the respective target tissue, with at least three transgenic embryos showing detectable signal in the target tissue (Figures 2B-D and S3). Moreover, four out of five synthetic heart enhancers are exclusively active in the heart, and one shows weak activity in the brain (Figures 2B and S3A). Similarly, four out of five synthetic CNS enhancers are CNS-specific, while one shows weak activity in the heart (Figures 2D and S3C). Interestingly, all five limb enhancers are strongly active in the limb and show weaker activity in other mesenchymal tissues expressing TWIST1^19^, i.e. craniofacial and trunk mesenchyme (Figures 2C and S3B).

## Discussion

Our findings establish that mammalian enhancer function is sufficiently encoded in DNA sequence to enable *de novo* design of tissue-specific enhancers that function *in vivo*. Whether reliable enhancer design would be feasible in mammals was unclear, given the structural and regulatory complexity of mammalian genomes, whose large size, extensive repeat content, CpG/dinucleotide biases, extensive sequence-, chromatin-, and epigenetic context dependence, and a large, partially overlapping TF repertoire could have obscured learnable rules. Despite these challenges, our results show that sequence-based models – anchored in chromatin accessibility and refined by transfer learning with limited sets of validated enhancers – can robustly guide enhancer design *in vivo*, as evidenced by reproducible activity of all fifteen synthetic enhancers in their intended tissues in the mouse embryo.

The success of the sequence-based design approach also yields insights into mammalian enhancer logic beyond model performance. Our results suggest that mammalian enhancer syntax is more systematic and learnable than often assumed, despite regulatory complexity. Furthermore, transfer learning might reflect a biologically meaningful decomposition: pre-training on genome-wide chromatin accessibility captures general regulatory context and broad regulatory cues, whereas fine-tuning on validated enhancers refines tissue-specific features, including key TF motifs. Finally, because the models distinguish functional enhancers from other accessible regions, inaccessible regions, and random synthetic sequences (Figures 1D and S1H), they likely captured *bona fide* sequence-level regulatory features rather than simply memorizing chromatin state.

Our study also shows that enhancer design in mammals does not require large, resource-intensive models or massive MPRA-scale datasets. Instead, compact CNNs, combined with genome-wide chromatin profiles that can be readily generated from any sample and with only a few hundred bona fide enhancers per tissue, are sufficient for robust design. As *in vivo* enhancer assays continue to scale^24,25^, our approach will be widely applicable to the design of more versatile and precise synthetic enhancers across mammalian tissues, cell types, and dynamic cell states.

### Scope of the study and outlook

This study focuses on the design of individual enhancer elements for a standardized reporter and promoter context. While this isolates enhancer-intrinsic regulatory logic and enables the design of synthetic enhancers sufficient for reporter expression in selected tissues, enhancer output in native loci can depend on promoter identity, chromatin environment, and higher-order genomic context. In addition, by focusing *locally* on individual elements, our approach does not address extended multi-enhancer gene regulatory architectures *globally* – which long-sequence genome models^26–30^ aim for – or cell types and developmental dynamics beyond the tissues and stage examined here. These limitations define the scope of this work and point to possible future extensions of the framework.

## Resource availability

### Lead contact

Further information and requests for resources and reagents should be directed to and will be fulfilled by the lead contact, Alexander Stark.

## Materials availability

This study did not generate new unique reagents.

## Data and code availability

The transcription factor motif database is available at https://www.vierstra.org/resources/motif_clustering. The weights and architecture of the final pre-trained sequence-to-accessibility and sequence-to-activity models are available at https://huggingface.co/Shenzhi-Chen/DeepSTARR-Mouse. The data used to train and evaluate the models are available at https://huggingface.co/datasets/Shenzhi-Chen/DeepSTARR-Mouse-dataset. In vivo enhancer activity data are publicly available on the VISTA Enhancer Browser website (https://enhancer.lbl.gov/vista/).

The code used to train the models and to make predictions on new sequences is available on GitHub (https://github.com/Shenzhi-Chen/DeepSTARR-Mouse). The code use for downstream analyses is available on GitHub (https://github.com/vloubiere/chen_loubiere_2025_git).

## Acknowledgements

We thank Bernardo P. de Almeida, Thomas Pierrot (InstaDeep) and Sasha Mendjan (IMBA) for discussions, Angela Andersen (Life Science Editors) for editorial support, and the members of the Chao Family Comprehensive Cancer Center Transgenic Mouse Facility and the IMP/IMBA/GMI IT department for their support. Research in the Stark group is supported by Boehringer Ingelheim GmbH, the Austrian Research Promotion Agency (FFG, FO999902549), the Austrian Science Fund (10.55776/P36971, 10.55776/PAT3564423) and the WWTF (10.47379/LS24012). This work was supported by National Institutes of Health grants DP2GM149555 and R01HD115268 (to E.Z.K.) and F30HD110233 (to E.W.H.). The Chao Family Comprehensive Cancer Center is supported in part by the National Cancer Institute of the NIH under award number P30CA062203. The content is solely the responsibility of the authors and does not necessarily represent the official views of the National Institutes of Health. For the purpose of Open Access, the author has applied a CC-BY public copyright license to any Author Accepted Manuscript version arising from this submission.

## Author contributions

S.C., V.L. and A.S. conceived the project. S.C. and V.L. performed all computational analyses and designed the synthetic enhancers. E.W.H., S.H.J., and A.D. performed mouse transgenic reporter experiments under the supervision of E.Z.K. S.C., V.L. and A.S. wrote the manuscript, with input from all authors. A.S. supervised the project.

## Supplemental information titles and legends

Document S1. Figures S1-S3.

Table S1. Designed synthetic enhancers selected for in vivo validation. Table S2. Publicly available ATAC-seq BAM files used in this study. Table S3. Full list of annotated ATAC-Seq peaks.

Table S4. Full list of VISTA Enhancer Browser sequences.

Table S5. Number of sequences used for sequence-to-accessibility models. Table S6. Number of sequences used for sequence-to-activity models.

## Methods

### Processing of ATAC-seq chromatin accessibility data

#### Peak calling

ATAC-seq mapped reads (mm10) from day 11.5 mouse embryos were obtained as BAM files from^14^ (see Table S2), and biological replicates were merged using samtools^31^ merge (v1.9). Peaks were called per biological replicate and on merged reads using MACS2^32^ (v2.2.5) with the following parameters: -g mm -f BAM --nomodel --shift -100 --extsize 200 --keep-dup all, and bedgraph pileup files were converted to bigwigs using the bedGraphToBigWig function^33^. Only confident peaks called in the two biological replicates and with merged reads were retained.

#### Peaks annotation

Confident ATAC-seq peaks from heart, limb, and CNS subregions (midbrain, forebrain, hindbrain, neural tube) were merged and resized to 1,001 bp (mean summit ±500 bp). Mean signal was computed from the corresponding merged bigwig files and log2-scaled as: log2(signal / sum[signal across all regions in that tissue] × 1e6). Outlier regions (scaled signal > 99.95th percentile) were removed and the remaining regions were clustered using a two-layer self-organizing map (kohonen R package v3.0.10^34^, supersom) on a 4×5 hexagonal, toroidal grid (layer 1: peak presence/absence per tissue (binary); layer 2: log2-scaled ATAC-seq signal). Peak clusters were further classified into 10 meta-clusters, including a cluster of “globally open” regions (accessible in all tissues, n = 17,212) as well as heart- limb- and midbrain-specific clusters (n = 3,008; 1,407 and 2,551, respectively). Weak clusters with no clear pattern were removed (n = 5,482). The full list of annotated peaks and the remaining regions (n = 40,196) is available in Table S3.

### Deep learning data preparation

#### VISTA sequences

In vivo enhancer activity data were downloaded from the VISTA Enhancer Browser^13^ on 01-09-2024. Sequences corresponding to mutated variants or with length >5 kb were removed, while sequences shorter than 1,501 bp were resized to a minimum length of 1,501 bp (n = 3,206; Table S4).

#### Inaccessible control regions

The mm10 version of the mouse genome was binned into contiguous 1,001bp bins, and bins located < 1.5kb away from the closest ATAC-seq peak or VISTA sequence were removed (for VISTA sequences of human origin, the mm10 coordinates of their orthologous regions were used, see Table S4). Then, 1 million bins were randomly sampled to serve as a pool of negative control regions.

#### Cross-validation scheme

The genomic coordinates of ATAC-seq peaks (n = 40,196), VISTA sequences (n = 3,206) and inaccessible control regions (n= 1,000,000) were combined and split into 20 strictly non-overlapping folds, ensuring a minimum gap of 1.5 kb between sequences assigned to different folds (for VISTA sequences of human origin, the mm10 coordinates of their orthologous regions were used, see Table S4). We then used a cross-validation setup where 18 folds were used for training, one for validation, and one for testing. However, regions on mouse chromosome 18 were systematically excluded from training and validation to serve as a fixed, “shared” test set across all models and tissues.

### Sequence-to-accessibility models

#### Data augmentation

All augmentations were applied after splitting regions into training/validation/test folds (see Cross-validation scheme). For each ATAC-seq peak and inaccessible control region, we included five 1,001-bp windows centered from −400 to +400 bp relative to the region center, with a 200-bp stride (−400, −200, 0, +200, +400). For peaks labelled as heart- limb- and midbrain-specific (see Table S3), we specifically reduced the stride to 50 bp in the corresponding tissue, resulting in a denser tiling (n = 17 windows per region). Finally, reverse-complemented sequences were added, doubling the number of sequences available for training.

#### Balancing

Inaccessible control regions were down-sampled to reach ∼1.5 times the total number of ATAC-Seq peaks after augmentation, to ensure reasonable class imbalances before training. The final number of sequences used for each tissue are available in Table S5. See data availability section to access the corresponding sequences.

#### Model architecture and training

For each region, DNA accessibility was computed from the corresponding merged bigwig file (containing the MACS2 treat pileup read coverage; see above) as log2(mean ATAC-seq signal + 1), and predicted using the previously described DeepSTARR CNN architecture^4^ with minor modifications. The CNN uses one-hot encoded 1,001 bp DNA sequences (A = [1,0,0,0], C = [0,1,0,0], G = [0,0,1,0], T = [0,0,0,1]) and four 1D convolutional layers (filters = 256,120,60,60; size = 7,3,3,3; padding = same), each followed by batch normalization, a ReLU non-linearity, and max-pooling (size = 3). The convolutional stack is followed by two fully connected layers (64 and 256 neurons), each followed by batch normalization, a ReLU non-linearity activation function, and dropout (fraction = 0.5). The final layer is mapped to the accessibility signal output. The models were implemented and trained in python (v3.7.15) using Keras (https://keras.io/, v2.4.3) from TensorFlow ^35^ (v2.4.1) using the Adam optimizer^36^ (learning rate = 0.005) with mean squared error as loss function, a batch size of 128, and early stopping with patience of five epochs.

To account for variance between different training runs, each model was trained in duplicates on three different held-out test folds, for a total of 6 models per tissue. For the three tissues, all models converged and showed neglectable variance in predictions, and predictions were therefore averaged across runs.

#### Model performance

Prediction bigwig tracks were generated for chromosome 18—which was excluded from all runs—using a 1,001 bp sliding window with a 200 bp stride. We then computed the Pearson correlation coefficient (PCC) between measured and predicted ATAC-seq signal over the union of heart, limb, and midbrain ATAC-seq peaks (Figure 1B). In addition, PCC was computed across the entire held-out test set for each tissue (Figure S1B) and on delta values between tissue pairs (Figure S1C), emphasizing performance at tissue-specific peaks. To compare models across tissues, we z-scored the observed DNA accessibility values across all held-out regions within each tissue. We then selected regions with a z-score > 1 in any tissue and clustered them using K-means (k = 6); and the resulting clusters were also used to classify the predicted values (Figure 1C). To compute the overlap between each K-means cluster and transcription start sites (TSSs) of mouse genes (Figure S1A), mm10 TSS coordinates were retrieved from GENCODE vM25.

#### Nucleotide contribution scores

We used DeepExplainer (the DeepSHAP implementation of DeepLIFT^37,38^; updated code at https://github.com/AvantiShri/shap/blob/master/shap/explainers/deep/deep_tf.py) to compute nucleotide contribution scores on the subset of held-out regions exactly centered on ATAC-seq peak summits (see Cross-validation scheme), using only the forward-strand sequence. For each input, 100 dinucleotide-shuffled versions were used as reference sequences, and final contribution scores were obtained by multiplying hypothetical importance scores by the one-hot–encoded matrix of the corresponding sequence. For each nucleotide, scores were averaged across all replicates and folds per tissue (n = 6).

### Sequence-to-activity models

#### Data augmentation

All augmentations were applied after splitting regions into training/validation/test folds (see Cross-validation scheme). VISTA sequences were tiled using a 1,001 bp sliding window with 50 bp stride. Only the windows with ≥950-bp overlap with the original region were retained and reverse-complemented sequences were added, doubling the number of sequences available for training. The final number of sequences used for each tissue are available in Table S6. See data availability section to access the corresponding sequences.

#### Model architecture and training

To classify DNA sequences based on their in vivo activity, we used a transfer-learning approach in which weights from the sequence-to-accessibility regression models were used to initialize a second CNN in the corresponding tissue ^5,15,16^. All layers were kept trainable, and the final layer’s activation was changed to a sigmoid function suitable for binary classification tasks. Models were trained with the Adam optimizer^36^ (with rate 1e-4), binary cross-entropy loss, a batch size of 128, and early stopping with patience of twenty epochs.

To account for variance between different training runs and improve the accuracy and robustness of the models, each model was trained in duplicates on three different held-out test folds, for a total of 6 models per tissue. For the three tissues, all models converged and showed neglectable variance in predictions, and predictions were therefore averaged across runs.

#### Model performance

For sequence-to-activity models after transfer learning, positive predictive value (PPV; TP/[TP

+ FP]) was computed on the held-out test set across increasing prediction thresholds (i.e., sequences were classified as positive when predicted activity ≥ threshold; Figure 1D) in each tissue, and we reported the maximum PPV achieved while ensuring that at least 100 predicted-positive sequences remained. We also evaluated two alternatives: (i) models trained directly on annotated VISTA enhancers without pre-training and (ii) predictions from the sequence-to-accessibility models (scaled to [0, 1] and used as activity predictions). In these cases, PPV was measured at the prediction thresholds corresponding to the maxima for the transfer-learning model in each tissue (Figures S1H–S1J).

#### Nucleotide contribution scores

Nucleotide contribution scores were computed with the approach described for the sequence-to-accessibility models, using the forward-strand sequences of all tiles within the held-out test set. For each nucleotide, scores were averaged across all replicates and folds per tissue (n = 6).

#### Design of tissue-specific synthetic enhancers

For each tissue, 1,200 random DNA sequences were generated with dinucleotide frequencies matched to VISTA sequences and used as input for Ledidi^22^ (v2.1.0), a model-guided gradient optimization approach. To limit the number and magnitude of edits, the edit-penalty parameter was set to 0.1. Seed sequences were then optimized using both the sequence-to-accessibility and sequence-to-activity models, with target predicted values set to 12 and 14, respectively. For activity optimization, the final sigmoid layer was removed, and pre-sigmoid logits were used to avoid gradient saturation and improve optimization stability. Designed sequences with a significant match to either the mouse or the human genome were identified using Blastn (via NIH NCBI Blast https://blast.ncbi.nlm.nih.gov/Blast.cgi) with default parameters (word size =11, Expect threshold= 0.05) and removed, yielding 1,049 usable sequences for heart, 1,020 for limb, and 978 for CNS/midbrain. From the designed sequences with high predicted activity in the target tissue (thresholds: >6 for heart, >5 for limb, >7 for CNS/midbrain) and low activity in the other two (<0; n = 654, 351, and 787 for heart, limb, and CNS/midbrain, respectively; see Figure S2C), fifteen were selected for in vivo validation (five per tissue). Selected sequences, their predicted activities and the corresponding nucleotide contribution scores are available in Table S1.

#### TF motifs analyses

Position weight matrices (PWMs) for transcription factor motifs listed in^39^ were mapped onto the held-out sequences using the matchMotifs function from the motifmatchr R package^40^ (v1.26.0; parameters: bg= ‘subject’, p.cutoff= 5e-05), and mean nucleotide contribution scores were computed for each motif instance; with respect to the sequence-to-accessibility or sequence-to-activity models. These per-instance means were then averaged per PWM (n = 2,174) and z-scored per tissue, and only PWMs with a z-score ≥ 2 for either accessibility or activity in any of the three tissues were retained (n = 358). Redundant PWMs were then filtered using the non-redundant motif clusters described in^39^, selecting only the PWM with the highest z-score as the representative of each cluster. Accessibility and activity z-scores were clipped at the 5th and 95th percentiles for each tissue, clustered with the hclust function in R (v4.4.1) using Euclidean distances, and the resulting tree was cut into four clusters with the cutree R function (Figure 2A).

#### Gene Ontology enrichment analysis

Mouse gene IDs associated with heart (GO:0007507), limb (GO:0060173), and central nervous system (GO:0007417) development were retrieved from the org.Mm.eg.db R package (v3.19.1). The list of ubiquitously expressed housekeeping genes was retrieved from^41^. For each of these four categories, the over-representation of the TFs associated to the four clusters defined in the previous section (n= 12, 55, 28, 56, see Figure 2A) was assessed using Fisher’s exact test, using all TFs represented in the PWM database as background (n= 599). p-values were corrected for multiple tested using the FDR method (significance threshold: FDR<0.05).

#### Single cell RNA-Seq analyses

Mouse single-cell RNA-seq data were retrieved from GSE119945, and only cells from day 11.5 embryos were considered (n = 602,722). We then selected the TF genes associated with the PWMs database described earlier (n = 573 after intersecting with the expression matrix), and cells not related to any of the target tissues (i.e., cardiac muscle lineages, limb mesenchyme, brain, or neural tube) were labeled as “other cell types”. Per-cell gene-level rankings were then computed on the raw count matrix using the AUCell R package (v1.26.0). AUC scores were finally computed for each of the TF clusters described earlier (Figure 2A) and for the remaining TFs (regrouped under the “Other TFs” label), before being z-scored and averaged per cell type for visualization (Figure S2B).

#### In vivo validation of synthetic enhancers

To test synthetic enhancer sequence activity in mouse embryos, each DNA sequence was custom ordered as a gBlock (IDT, San Diego, CA) (see Table S1 for sequences). Synthetic enhancers were then inserted into a PCR4-Shh::lacZ-H11 vector backbone (Addgene #139098) using Gibson-based assembly methods (Gibson et al, Nat Methods 2009). Sequence identity was confirmed via enzymatic digestion and Sanger sequencing. A mix of (i) Cas9 protein (final concentration of 20 ng/μl; IDT Cat. No. 1074181), (ii) sgRNA (50 ng/μl), and (iii) donor plasmid (7 ng/μl) was prepared in injection buffer (10 mM Tris, pH 7.5; 0.1 mM EDTA) and injected into the pronucleus of FVB embryos. Mice (FVB and CD-1 strains) were kept in standard housing conditions (temperature 19–23 °C and humidity 40–60%) with food and water provided ad libitum on a reversed 12-h dark–light cycle. Pregnant dams were humanely euthanized, and E11.5 embryos were carefully removed under bright-field stereoscopes in ice-cold PBS (Cytiva, SH30256.01). Both sexes of embryos were presumed to be included. Yolk sacs were collected for genotyping, and successful integration events at the H11 locus were determined by PCR using primers described previously^23^. Scoring of tissue positivity was performed for each embryo by two independent scorers in a non-blinded fashion. This research complies with all relevant ethical regulations. All animal procedures, including those related to generating transgenic mice, were conducted in accordance with the guidelines of the National Institutes of Health (NIH) and approved by the Institutional Animal Care and Use Committee at the University of California, Irvine under protocol no. AUP-23-005.

## Quantification and statistical analysis

All statistical analysis and plots were done in R^42^ (v4.4.1) using the data.table package^43^.

