## Supplementary material for "Predictive design of tissue-specific mammalian enhancers that function *in vivo* in the mouse embryo": Document_S1

### Figure S1

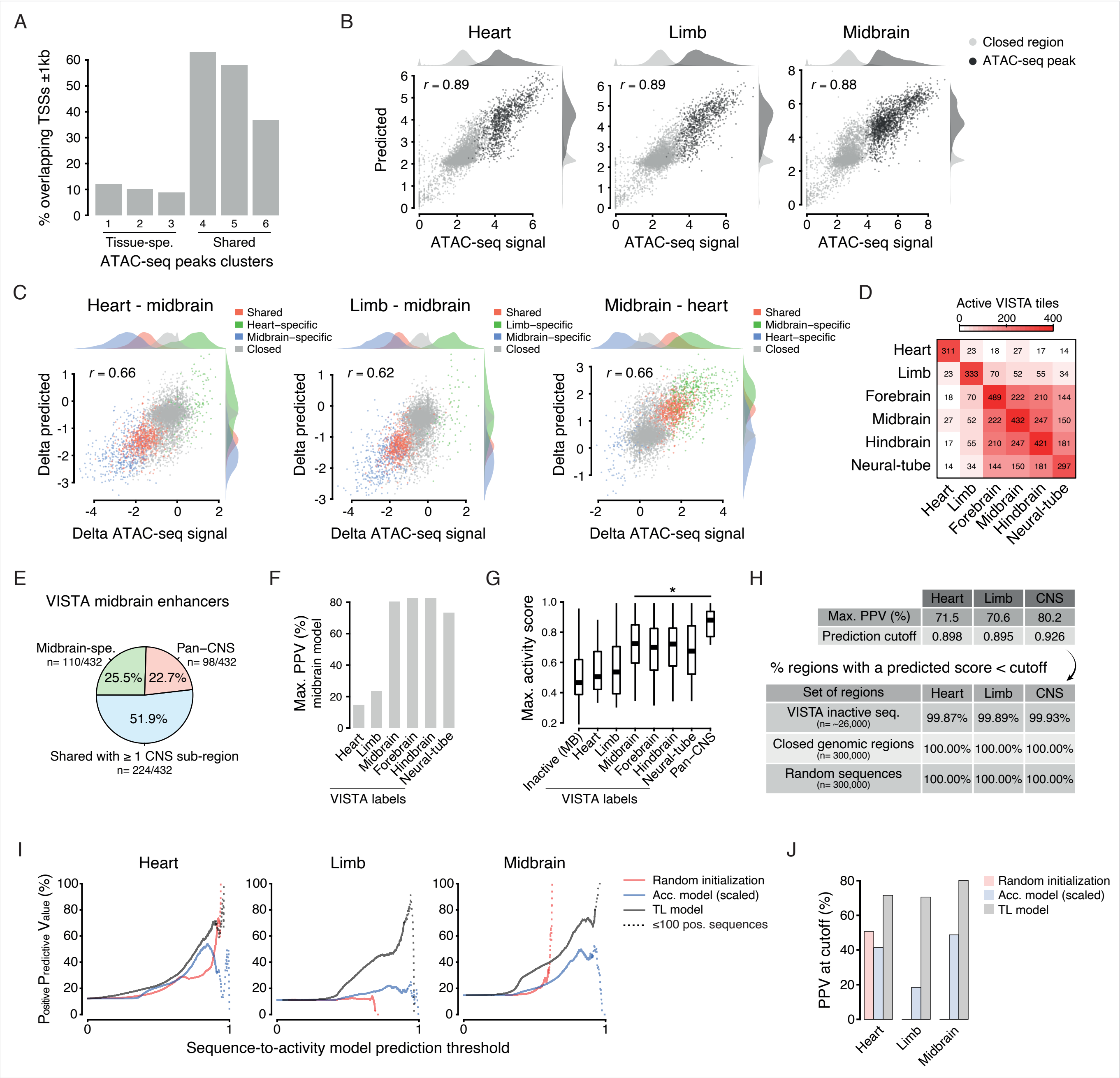


**Figure S1. Evaluation of sequence-to-accessibility and sequence-to-activity models, related to Figure 1**

(A) Percentage of ATAC-seq peaks overlapping Transcription Start Sites (TSSs) +/- 1kb (for each cluster defined in Figure 1C).

(B) Observed vs. predicted ATAC-seq signal on sequences that were not used for training (test set).

(C) Delta observed vs. predicted ATAC-seq signal on sequences that were not used for training (test set), further stratified based on their tissue-specificity (color legend).

(D) Overlap between tissue labels associated to validated enhancer sequences from the VISTA enhancer browser.

(E) Pie chart indicating the percentage of VISTA enhancers active in the midbrain that are: specifically active in the midbrain (green), active in at least one other CNS subregion (blue), or active in all CNS subregions (red).

(F) Maximum PPV (%) achieved by the midbrain sequence-to-activity model, evaluated on VISTA enhancers active in heart, limb, or any of the four CNS subregions (x-axis). As in Figure 1D, the maximum PPV is reported when at least 100 positive sequences remain.

(G) Activity score for VISTA sequences that are inactive in midbrain (MB) versus those active in heart, limb, any of the four CNS subregions, or all CNS subregions (pan-CNS; x-axis). For each VISTA sequence, only the maximum predicted activity score across its tiles is reported.

(H) Top: for each tissue, the maximum PPV (%) achieved by the respective model is shown, together with the corresponding prediction threshold. Bottom: percentage of sequences predicted to be inactive (predicted activity score < prediction cutoff defined in the top panel), for three different sets of held-out control regions.

(I) PPV of enhancer activity across prediction score thresholds, comparing a scaled sequence-to-accessibility model (blue) and a randomly initialized model trained directly on VISTA enhancers (red). The sequence-to-activity model obtained by transfer learning – introduced in Figure 1D – is shown as a reference (TL, black). For each threshold (x-axis, 0–1), the y-axis shows the percentage of active sequences among predicted positives (predicted score ≥ threshold).

(J) PPV achieved at the prediction cutoffs defined in panel H, for the three approaches described in panel I (see color legend).

### Figure S2

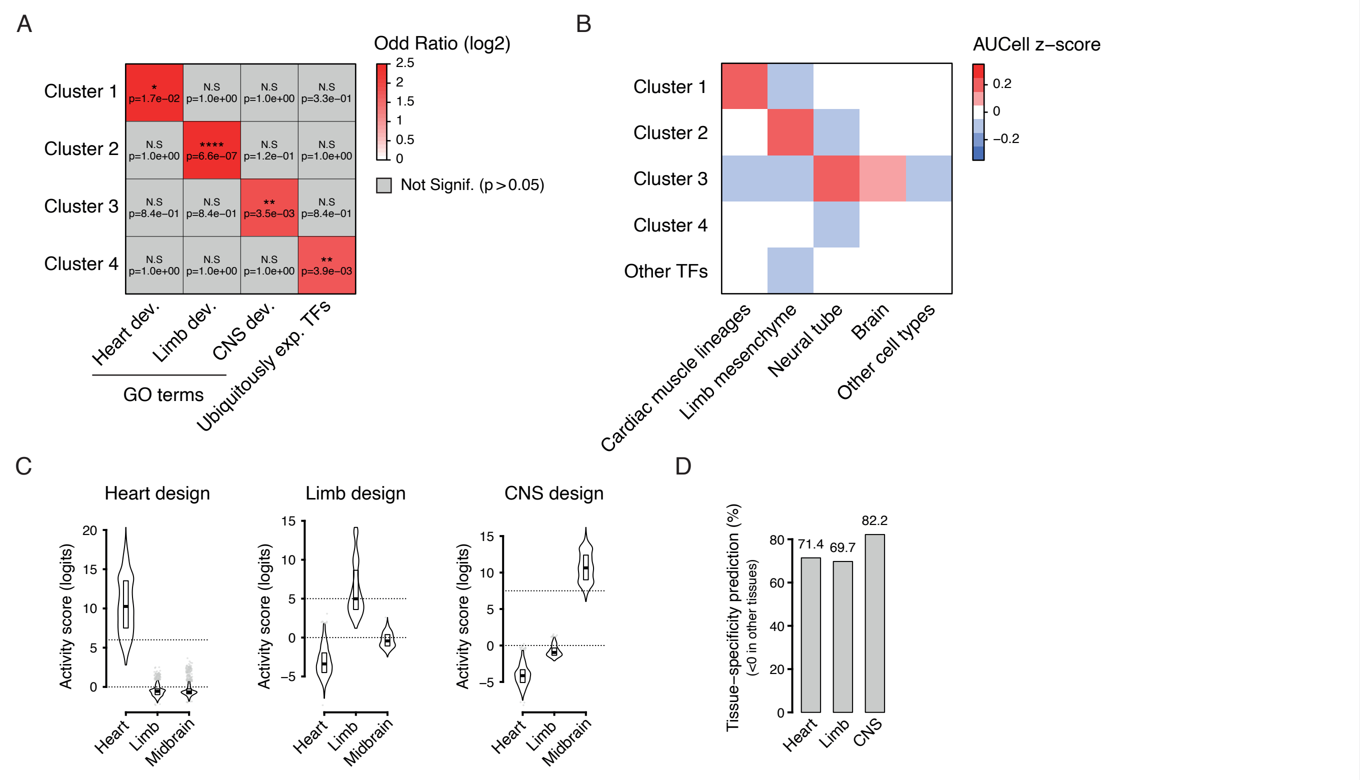


**Figure S2. Evaluation of tissue specificity at the level of TF motif content and enhancer-activity prediction, related to Figure 2**

(A) For each motif cluster defined in Figure 2A, the corresponding TFs were retrieved, and their over-representation across GO categories (heart development, limb development, CNS development) and ubiquitously expressed TF genes (housekeeping) was assessed using Fisher's exact test. The color legend indicates the log2(odds ratio), while P values adjusted for multiple testing are shown in plain text.

(B) Single-cell RNA-seq–derived AUCell z-scores for TFs corresponding to each TF motif cluster defined in Figure 2A (see Methods).

(C) Predicted activity scores (logits) for synthetic enhancer sequences designed for heart (left), limb (center), or CNS (right), using the three sequence-to-activity models (x-axes). The dotted lines indicate the cutoffs used to define sequences predicted to be strongly active in the target tissue (>6, >5, and >7 for heart, limb, and midbrain, respectively) and inactive in the two other tissues (<0).

(D) For sequences predicted to be strongly active in their target tissue (using the cutoffs defined in panel C), percentage of sequences that were predicted to be inactive (<0) in the two other tissues.

### Figure S3

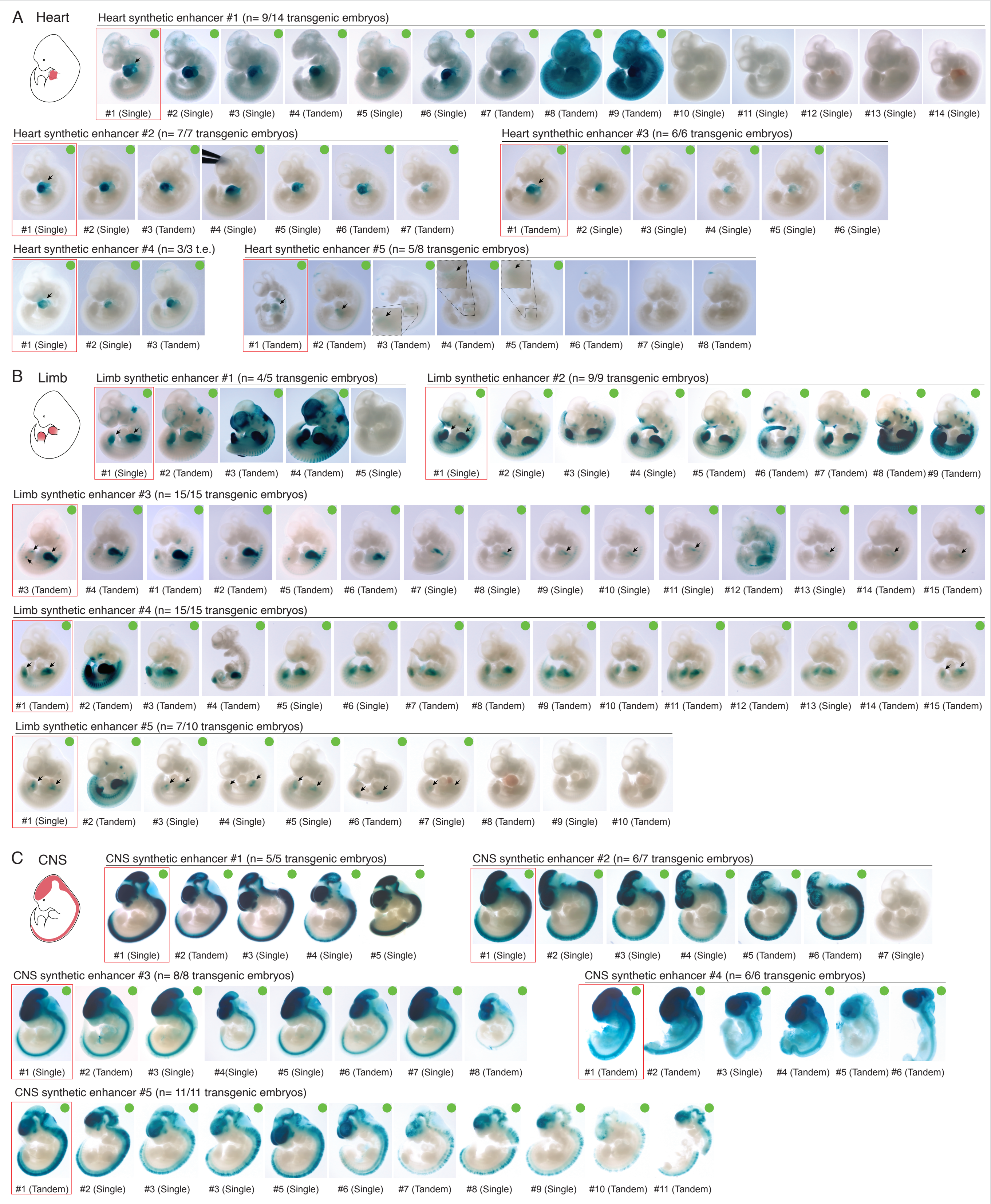


**Figure S3. Overview of all transgenic mouse embryos for enhancer validation, related to Figure 2**
(A) On the top left, a schematic view of E11.5 mouse embryos highlighting the position of the heart (in red) is shown. For each of the five heart synthetic enhancers that were tested in vivo, all correspond lacZ-stained E11.5 mouse embryos are shown. Following VISTA convention[13], a sequence was considered reproducibly active in the target tissue if at least three embryos showed detectable staining in that tissue. Positive embryos are highlighted using a green circle on the top right part of the image. Red boxes indicate the representative embryos that are shown in Figures 2B-2D and are reproduced here for completeness.
(B-C) Same as panel A but for limb and heart tissues (n= 5 synthetic enhancers per tissue).
